# Integration of QTL Mapping, Transcriptomics, and Genome Resequencing Identifies Yield-Associated Genes for Salt Stress in Rice

**DOI:** 10.64898/2026.03.31.715716

**Authors:** Nivesh Kumar, Bikram P Singh, Pragya Mishra, Mukta Rani, Anoop Gurjar, Ayushi Mishra, Anshuman Shah, Nitin Gadol, Sushma Tiwari, Suman Rathor, P C Sharma, SL Krishnamurthy, Teruhiro Takabe, Shiro Mitsuya, Sanjay Kalia, Nagendra K Singh, Vandna Rai

**Author notes:** Corresponding author: Vandna Rai.

## Abstract

Salinity and sodicity stresses adversely affect rice growth and yield. To overcome yield losses, suitable tolerant rice cultivars can be developed through a marker-assisted breeding (MAB) program. In the present study, genomic regions associated with sodicity stress tolerance at the reproductive stage were identified using a high-density 50kSNP array in a recombinant inbred line (RIL) population derived from the contrasting rice genotypes CSR11 and MI48. A total of 50 QTLs were detected for various yield-related traits; further, 19 QTLs with ≥15% of phenotypic variance were selected for integrated (omics) analysis. RNA sequencing of leaves and panicles at the reproductive stage under sodic stress conditions was employed to find differentially expressed genes. A total of 1368 and 1410 SNPs; 104 and 144 indels were found for MI48 and CSR11, respectively, within the QTL regions from resequencing. At chromosomes 1 and 6, colocalized QTLs (*qPH1-1/qGP1-1* and *qGP6-2*/*qSSI6-2*) were discovered. Differentially expressed genes (DEGs) were mapped over the QTL regions selected, and SNP variations and indels were screened for colocalized QTLs. Potential candidate genes, namely Os-pGlcT1 (Os01g0133400), OsHKT2;1 (Os06g0701600) and OsHKT2;4 (Os06g0701700), OsANTH12 (Os06g0699800), and OsPTR2 (Os06g0706400), were identified as being responsible for glucose transport, ion homeostasis, pollen germination, and nitrogen use efficiency, respectively, under salt stress. Finally, our study provides important insights into the genes and potential mechanisms affecting grain yield under sodic stress in rice, which will contribute to the development of molecular markers for rice breeding programs.

## Introduction

Rice is the staple food crop for over half of the world’s population (∼3.5 billion), and demand for rice is increasing due to population growth; therefore, production must increase by 2050 to feed a population of 9.8 billion (Contreras-Moreira et al., 2026). The crop is cultivated under rain-fed conditions in various agro-climatic zones and subjected to various abiotic stresses, including salinity (Singh et al., 2016).

More than 1 billion ha of land are affected by salinity, of which approximately 60% are affected by sodicity (FAO 2024). The sodic soil is dominated by sodium bicarbonate and carbonate salts. Due to global warming and freshwater scarcity, more than 50% of land will be salt-affected by 2025 (Zhang et al., 2023). Salt stress triggers an array of morpho-physiological and molecular responses in rice that adversely affect growth, development, and yield potential (Munns and Tester, 2008; Krishnamurthy et al., 2016; Ismail and Horie, 2017; Krishnamurthy et al., 2023; Krishnamurthy et al., 2024; Rathor et al., 2026). Therefore, global efforts have been made using either conventional or biotechnological interventions to engineer traits to alleviate salinity stress responses (Gregorio 1997; Deinlein et al., 2014; Mickelbart et al., 2015). One of the conventional approaches has been the identification of quantitative trait loci (QTLs), which link genetic markers such as amplified fragment length polymorphism (AFLP), restriction fragment length polymorphism (RFLP), and microsatellites. Saltol and other QTLs for salinity tolerance for the seedling stage were reported in different studies (Gregorio 1997) using markers. Subsequently, sequencing of the rice genome (IRGSP, 2005) has provided a fillip towards the identification of QTLs using simple sequence repeats (SSRs), single-nucleotide polymorphic (SNPs) markers and genome-wide association studies (Kumar et al., 2025; Tiwari et al., 2016; Mishra et al., 2016; Warraich et al., 2021; Rathor et al., 2022). Through QTLs, regions responsible for salt tolerance can be identified, but reaching the gene level is still a daunting task.

Salinity stress tolerance in plants is a polygenic trait (Lin et al. 2004). Several molecular entities (TFs, transporters, etc.) have been identified as playing crucial roles in sensing and signalling cascades governing salinity stress responses in glycophytes (Chinnusamy et al. 2006; Kumari et al. 2009; Mishra et al. 2016). Thomson et al. (2010) identified distinct Pokkali alleles in the Saltol region and highlighted the possibility that the sodium transporter gene SKC1, located at 11.46 Mb and first reported in Nona Bokra (Ren et al. 2005), may confer salinity tolerance at the seedling stage. Efforts have also been made to manipulate many of these molecular entities, like Na^+^ transporters (HKT1;4, HKT1;5 NHX) (Suzuki et al., 2016; Kobayashi et al., 2017), ROS scavengers (Lou et al., 2018), serine hydroxymethyltransferase3 (Mishra et al., 2019), trehalose (Joshi et al., 2020) and kinases like CIPK (calcineurin B-like protein-interacting protein) (Xiang et al., 2007) to engineer rice that could sustain growth under salinity stress.

A linkage map with high-density molecular markers is a prerequisite for successful QTL identification (Hackett et al., 2013). Amongst all types of molecular markers, SNPs have proven to be the marker of choice because of their ubiquitous presence in high numbers, uniform distribution, bi-allelic nature and high heritability (Mammadov et al., 2012). Therefore, identifying SNPs linked to important agronomical traits is of utmost importance. The availability of high-density SNP chip arrays enables high-throughput genotyping of large populations within a short period. A 50K- and 90K-SNP chip was designed and validated for high-throughput genotyping in rice (Singh et al., 2015; Daware et al., 2023). Progress in next-generation sequencing techniques has generated large-scale genome and transcriptome sequencing data to better understand salinity stress responses in *O. sativa* and wild rice species under salt stress conditions (Mizuno et al., 2010; Garg et al., 2014; Mondal et al., 2017; Razzaque et al., 2017). Gene expression studies, in conjunction with genetic mapping, were also used to identify gene expression within the QTL region for salinity stress responses. Bulk segregant assay (BSA) with recombinant inbred lines (RILs) developed by salt-tolerant (CSR27) and salt-sensitive (MI48) was employed, and genes (an integral transmembrane protein DUF6 and a cation chloride cotransporter) were found to be co-located in the QTL interval (Pandit et al., 2010). Similarly, RNA sequencing was used to assess cold-responsive gene expression, and the genome-wide density of differentially expressed genes was linked to QTLs for the seedling stage of rice (Buti et al., 2018). All these techniques have been used for seedling-stage salt stress tolerance in rice. Using a RIL population of Liangxiang5 and 03GY28, *qLeafColor9.1*, a major QTL, was found on chromosome 9, wherein *OsSGR*, a senescence-related gene, showed higher expression under salt stress (Zhang et al., 2026). In another study, genomic and transcriptomic data were integrated to identify introgressed regions and differentially expressed genes in the introgression line JN100 derived from Nona Bokra and Jupiter (Chaudhary et al., 2025), and the genes primarily belong to the CHXs and NHXs transporter families and transcription factors.

Rice is sensitive during the seedling and reproductive stages of development. However, there is no correlation between the two stages in response to salinity, indicating distinct mechanisms of salt stress tolerance (Moradi et al., 2003). Salt stress reduces grain yield at the reproductive stage, leading to panicle formation, spikelet initiation, pollen grain germination, floret fertilization, and, therefore, sterile florets (Lauchli and Grattan, 2007). The genetic basis of salt stress tolerance was analyzed in the RIL population developed from Pusa44 and Bura Rata, and a positive correlation was observed between grain yield and spikelet fertility, leading to the identification of some RILs with ≤75% spikelet fertility (Rathore et al., 2026). Saltol QTL identified is conferring tolerance to salinity at the seedling stage, but does not confer tolerance at the reproductive stage. Thus, identifying QTLs/candidate genes and understanding the salinity-sodicity tolerance mechanism at the reproductive stage remained challenging. The present study aims to identify genomic regions that contribute to salt stress tolerance in rice at the reproductive stage. In the present study, an array of SNPs representing high-density linkage maps, parental resequencing, and whole-genome transcriptome profiling was used to identify candidate genes and elucidate the molecular mechanisms underlying salt stress tolerance in tolerant and sensitive rice cultivars.

## Materials and Methods

### Plant Material, Phenotyping for Salt Stress Tolerance and QTL Mapping Populations

The plant material used in the present study is the RIL population derived from CSR 11 (salt-tolerant) and MI48 (salt-sensitive) varieties. Phenotyping was conducted on parents and RIL populations for salt tolerance parameters over three years (Tiwari et al., 2016). Phenotyping was done under normal (pH ∼ 7.5), moderate (pH ∼ 9.5) and high (pH ∼ 9.8) sodic stress conditions in a controlled microplot. Parameters recorded for phenotyping were Panicle Length (PL), Productive Tillers per plant (P), Total Tillers per plant (TT), 1000 Seed Weight (SW), Plant Height (PH), Spikelet Fertility (SF), Grain Yield (GY), Days to fifty percent Flowering (DTF) and Grains per Panicle (GP), and also the Stress Susceptibility Index (SSI) calculated from the grain yield.

### Genotyping by 50K SNP Chip

The methodology described in Tiwari et al. (2016) was followed for genotyping, except that, instead of pooling for bulked segregant analysis, each of the 96 RILs in the CSR11/MI-48 mapping population was used for genomic DNA extraction and SNP genotyping. The SNP genotyping was performed using the 50K SNP chip (OsSNPnks) on the GeneTitan instrument, which provided the allele calls for all genotypes (Singh et al., 2018). The data quality was measured by DQC (development quality check) and QCR (quality call rate) tests. DQC measures the extent to which the distribution of signal value is resolved from the background value. DQC value 0 indicated no resolution, while 1 indicated perfect resolution. The recommended value of DQC is ≥0.82. QC call rate was the fraction of called SNPs per sample over the total number of SNPs in a dataset. The threshold value for the QC call rate was ≥94%.

### High-Density Genetic Linkage Map Construction

50051 SNP markers encrypted on the chip were used to analyze the samples. Passed samples based on DQC versus QC call rate tests were selected, and their genotype data were used to construct the linkage map. A genetic linkage map was constructed from the allele calls for RIL’s by using the JoinMap 4.1 program (Van Ooijen et al., 2006). The software processed the genotypic data to remove monomorphic markers, duplicate markers, segregation-distorted markers, and markers missing in either parent or in more than the given threshold number of individuals. Processed polymorphic markers were used for linkage map construction, with LOD values≥3 and recombination frequencies ≤0.25. The recombination frequencies were converted to centimorgans (cM) using the Kosambi map function (Kosambi, 1943). The final marker positions for each chromosome were visualized using MapChart 2.2 (Voorrips, 2002).QTL mapping was done by WinQTLCart 2.5 software. The five input files were created: chromosome labels and several marker information files, marker labels, marker positions, genotype data, and phenotype data. All five input files were loaded according to the manual instructions, and QTL were detected using 1000 bootstrap replicates at a P-value of 0.05.

### RNA Sequencing

To observe transcriptional changes in rice induced by salt stress, the CSR11 and MI48 parental sets were grown in controlled sodic microplots. The panicles and leaves of tolerant and sensitive plants were harvested in replication at the reproductive stage for transcriptome analysis. The stored tissues of individual leaves and panicles were then used for total RNA extraction from CSR11 and MI48 genotypes using the Spectrum Plant Total RNA kit (Sigma), followed by the manufacturer’s instructions. To remove residual genomic DNA, the RNA samples were purified with DNase. Later, the RNA integrity was confirmed by running 1% formaldehyde agarose gel. The purified RNA was then used for transcriptome sequencing.

### Annotation and differential gene expression analysis

Raw sequences in FASTQ format obtained from the Illumina HiSeq platform were analyzed using the Fastp tool. The sequences were aligned and mapped to the IRGSP-1.0 reference genome sequence using HISAT2. The expression level of each transcript was reported as fragments per kilobase per million mapped reads (FPKM), calculated from the number of mapped reads. EdgeR was used to detect differentially expressed genes from at least two replicates, with a correlation coefficient>0.90 across libraries based on FPKM values. Pathway and gene ontology analyses were performed using PlantCyc, Reactome, and Ensembl. To identify DEGs, the threshold was applied as (log2 (fold change)) >2.0 or <-1 and p-value <0.05. For instance, the Venny software (http://bioinfogp.cnb.csic.es/tools/venny/index2.0.2.html) was used to construct Venn diagrams to represent DEGs in the leaf and panicle. Moreover, it was used to demonstrate the list of genes that shared a unique expression profile under salt stress conditions.

### Resequencing

The parental genotypes were resequenced, and the raw data were subjected to quality control to assess data quality. The raw data were found to be of good quality and were processed for alignment against the high-quality reference genome. The aligned reads were sorted, and putative duplicates were removed, followed by variant calling and variant annotation. The raw variants were filtered at a read depth cutoff of 5 to proceed with the downstream analysis, such as QTL identification. The raw data were processed using the NGS QC Toolkit (Patel and Jain, 2012) to assess the quality of the sequenced data. The PE 2 X 150 sequenced data was generated with 10x Coverage. The alignment was performed using Bowtie2 version 2.4.1 (Langmead and Salzberg, 2012). against the rice reference genome, and the alignment percentage of the reads with the reference genome was found to be good. The aligned reads were considered for downstream variant calling. The variant calling was performed using samtools version 0.1.19 (Li, 2011). Variants were filtered at a read depth >=5 to exclude SNPs and InDels identified at a read depth cutoff of 5.

## Results

### Phenotypic Analysis for Salt Stress Tolerance

The 216 RILs derived from the CSR11/MI48 parent population were evaluated phenotypically over three consecutive years. The phenotypic data indicated significant variation in the nine agronomic characters evaluated under moderate (pH ∼ 8.5) and high (pH ∼ 9.8) sodic stress environments, thus rendering it a suitable population for QTL analysis (Table 1). Moreover, the frequency distribution for phenotypic values remained continuous with normal or skew-normal distributions. Besides, these traits conferred the transgressive segregation in the RILs of CSR11/MI48, implying that alleles with positive effects on certain traits were distributed among the parents. Furthermore, the correlation coefficient indicated that these traits were significantly and positively correlated across all three seasons (Tiwari et al., 2016).

**Table 1.**
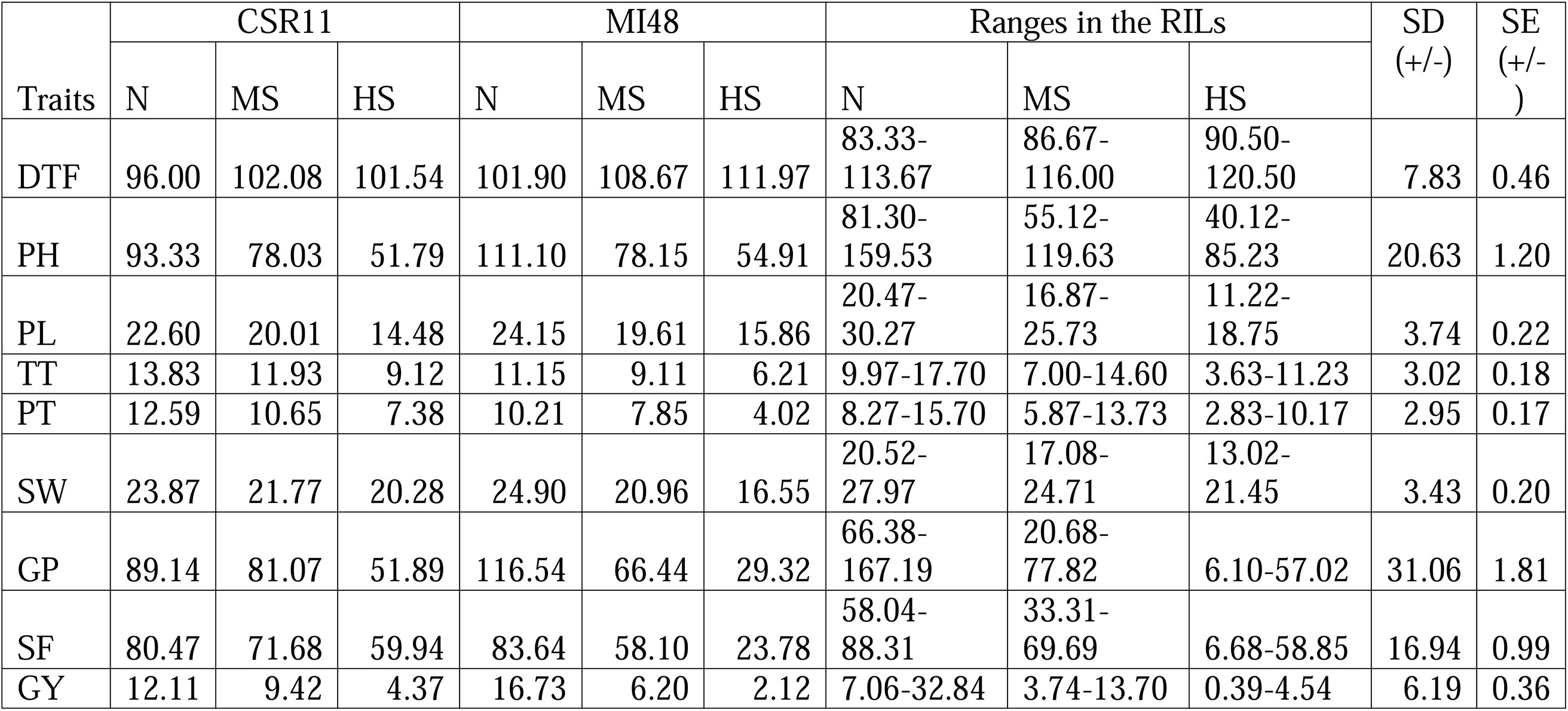
Variation for yield and yield contributing traits among 96 RILs derived from CSR11/MI48 over three seasons under control (N), moderate sodic (MS) and high sodic (HS) conditions.

### A High-Density Genetic Linkage Map for Sodic Stress Tolerance

RILs (96) derived from the CSR11×MI48 crosses were randomly selected to construct data for sodic stress tolerance and analyze a genetic linkage map. It revealed that a total of 12147 markers were polymorphic in these RILs with DQC thresholds (>0.85) and SNP call rates (>95%). The Affymetrix Genotyping Console™ v4.1 software package was used to extract polymorphic markers for the genotyping data analysis. Furthermore, to narrow the marker distribution, only one SNP per gene was selected from the 12147 polymorphic SNP markers. As a result, 1401 SNP markers were segregated in RILs and parents with statistically significant chi-square values. The linkage map contained 1401 markers spanning 1753.77 cM across the 12 rice chromosomes, as shown in (Supplementary Fig 1. Moreover, the total length of Chr 2 is 208.46 cM, the largest, and Chr 4 is 83.93 cM, the smallest. The number of markers per linkage group ranged from 206 (Chr 2) to 34 (Chr 10), with an average of 116.35 markers per chromosome. A summary of the constructed genetic map is presented in Table 2.

**Table 2:**
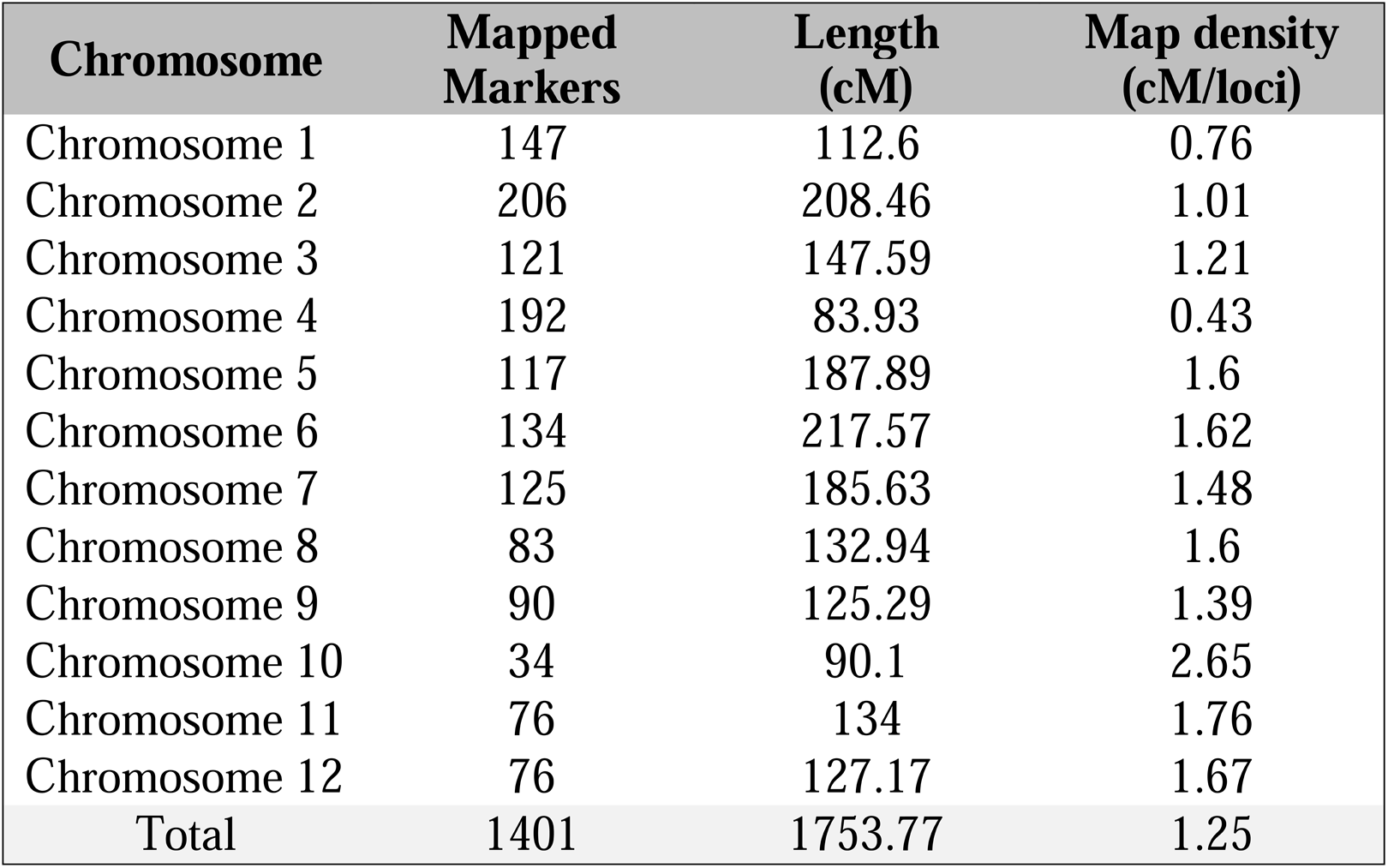
Summary statistics of the rice genetic linkage map constructed using the RIL population derived from a cross between (CSR11 X M148)

### Identification of QTLs for Sodic Stress Tolerance

Genotyping data from 1401 markers were analyzed alongside phenotyping data on sodic stress-related parameters over three consecutive years to map QTLs involved in salt stress tolerance. With the application of composite interval mapping (CIM) in Windows QTL Cartographer 2.5, a total of 50 QTLs were identified for the following traits: PL, PT, TT, SW, PH, SF, GY, DTF, GP and SSI. They were distributed throughout the rice genome except chromosome 10 (Table 3, Supplementary Fig. 1). The LOD thresholds ranged from 3.12 (*qPL6-2*) to 7.05 (*qGP1-1*). The identified QTLs with PV ≥15% were summarized in Table 3 and plotted for each locus on the linkage map (Supplementary Fig. 1). The identification of QTLs associated with specific parameters in 96 RILs derived from the CSR11 X M148 mapping population over the three consecutive years is explained here.

**Table 3.**
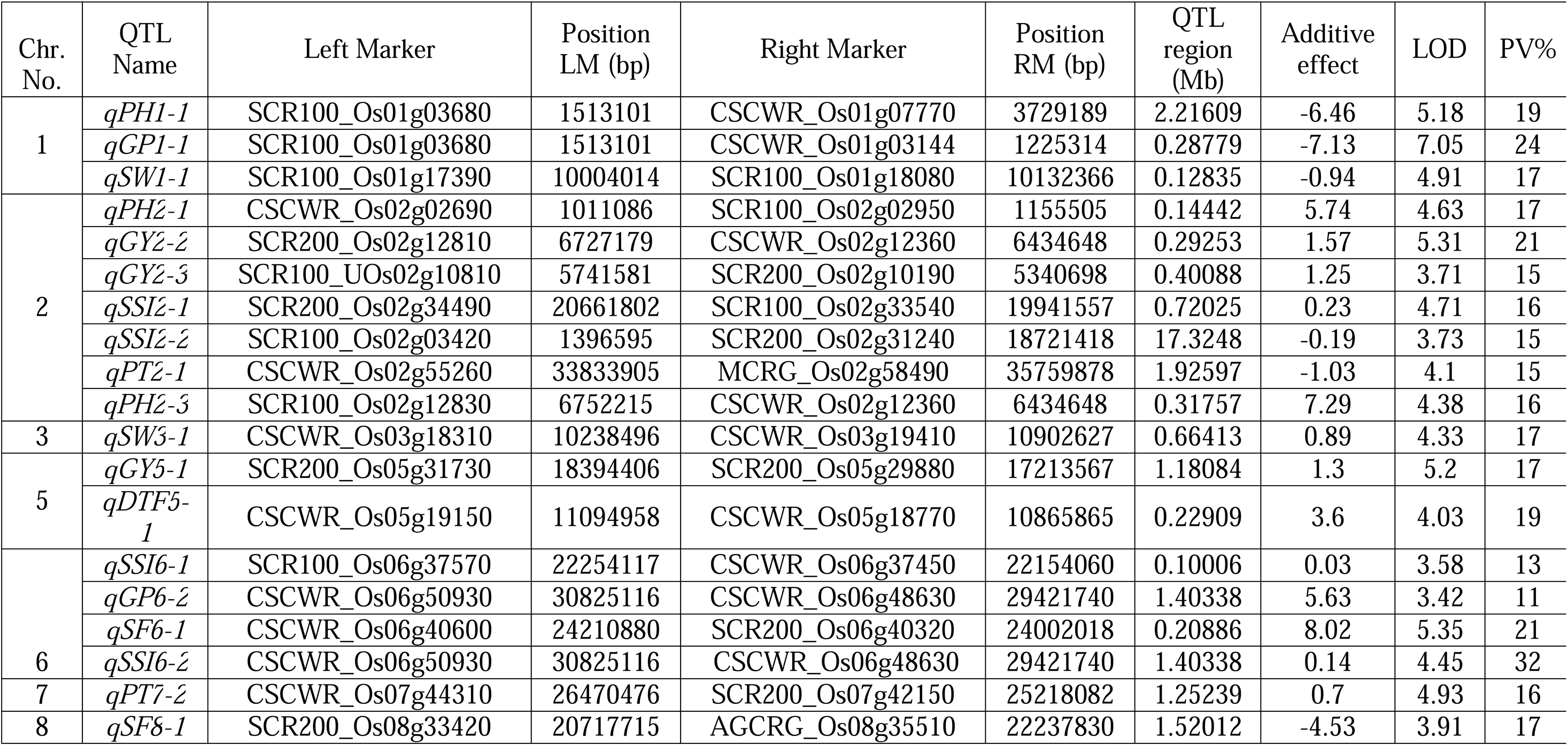
Quantitative trait loci (QTLs) associated with the traits: Panicle Length (PL), Productive Tillers per plant (PT), Total Tillers per plant (TT), 1000 Seed Weight (SW), Plant Height (PH), Spikelet Fertility (SF), Grain Yield (GY), Days to fifty percent Flowering (DTF), Grains per Panicle (GP), and Stress Susceptibility Index (SSI) and mapped on the linkage map of rice (CSR11XMI48). A negative value indicates that given trait is derived from MI48 and a positive number indicates that the trait is derived from CSR11.

Over the three years, a significant variation was observed in the panicle length of the tolerant RILs compared with controls (without stress). Six QTLs for PL were identified within the mapping population, among which four were mapped on chromosome 2 in years 1, 2 and 3, and two were mapped on chromosome 6 in years 1 and 2. They accounted for an estimated 12% to 14% of the phenotypic variance explained (PVE) and LOD thresholds ranging between 3.12 (*qPL6-2*) to 4.20 (*qPL2-4*). A single major QTL (*qDTF5-1*) was detected for DTF on chromosome 5 in year 3, accounting for 19% of PVE and 4.03 LOD. CSR11 contributed the positive allele for the *qDTF5-1* under the high sodic stress. Four QTLs were identified for GP within this RIL mapping population, of which one mapped to chromosome 1 in year 1 and another to chromosome 4 in year 3. Two QTLs were mapped on chromosome 6 in years 1 and 2, respectively. They accounted for an estimated 11% to 24% of the PVE and LOD thresholds, ranging between 3.42 (*qGP6-2*) and 7.05 (*qGP1-1*). Interestingly, a substantial phenotypic variation was observed among the RILs for grains per panicle under medium-sodicity tolerance in year 1.

A total of seven QTLs were identified for GY, among which three QTLs were mapped on chromosome 2 in year 2, whereas single QTLs were identified on chromosomes 4, 5 and 12 in year 1 and single QTLs on chromosome 7 in year 2. They accounted for an estimated 9% to 21% of the PVE and LOD thresholds, ranging between 3.68 (*qGY2-1*) and 5.31 (*qGY2-2*). MI48 contributed the negative allele for *qGY4-1* under the medium sodicity, whereas CSR11 contributed the other six favourable alleles. Six QTLs were identified for PH, among which two were mapped on chromosomes two and 4 in years 2 and 3, whereas single QTLs were mapped on chromosomes 1 and 7 in years 1 and 3, respectively. They accounted for an estimated 12% to 19% of the PVE and LOD thresholds, ranging between 3.58 (*qPH4-1*) and 5.18 (*qPH1-1*). Interestingly, one major QTL (*qPH1-1*) derived from the MI48 parent, accounting for 19% of the phenotypic variance, was mapped near the previously reported plant height green revolution gene Sd1, *LOC_Os01g6610* (Spielmeyer et al., 2002).

Five QTLs were identified for PT; three were mapped on chromosome 7 in years 1 and 2, whereas single QTLs were mapped on chromosomes 5 and 2 in years 2 and 3. They accounted for an estimated 11% to 18% of PVE and LOD thresholds ranging between 3.21 (*qPT5-1*) to 4.93 (*qPT7-2*). Nine QTLs were identified for SF, among which two were mapped on chromosomes 7 and 9 in years 1 and 3, whereas single QTLs were mapped on chromosomes 4 in year 1 and 6, 12 in year 2 and on the same 8 and 2 in year 3. They accounted for an estimated 11% to 22% of PVE and LOD thresholds ranging between 3.44 (*qSF12-1*) to 5.35 (*qSF6-1*).

Six QTLs were identified for SSI, among which two were mapped on chromosomes 2 and 6 in all three consecutive years, whereas single QTLs were mapped on chromosomes 5 and 1 in year 1 and 3, respectively. They accounted for an estimated 11% to 32% of PVE and LOD thresholds ranging between 3.23 (*qSSI1-1*) to 4.71 (*qSSI2-1*). Moreover, a major QTL (*qSSI6-2*) on chromosome 6 accounted for an estimated 32% of PVE and a LOD threshold score of 4.45. Tiwari et al. (2016) identified 21 QTLs for SSI using a bulk segregation analysis approach with the same SNP chip. There were three QTLs on chromosomes 1 and 2 and five on chromosomes 5 and 6. Our findings showed that the QTLs reported on these chromosomes were consistent with both approaches, BSA and traditional mapping. Also, the previously reported QTL *qSSIGY6.4* was found near *qSSI6-1* reported in this study.

Four QTLs were identified for SW, among which single QTLs were mapped on chromosomes 8, 11, 1 and 3 in years 1, 2 and 3. They accounted for an estimated 13% to 17% of PVE and LOD thresholds ranging between 3.36 (*qSW8-1*) and 4.91 (*qSW1-1*). The major QTL *qSW1-1* was localized near the *qSKC1* region and the previously reported QTLs *qNa/KSH-1.1* and *qKSH*, which were reported on the CSR27 and MI48 mapping populations by Pandit et al. (2010). Two QTLs were identified for TT and mapped to chromosomes 6 and 7 in years 2 and 3, respectively. They accounted for an estimated 12% and 13% of PVE, respectively, and LOD thresholds scored 3.64 (*qTT6-1*) to 3.68 (*qTT7-1*).

### Whole genome resequencing of contrasting parents

To identify genetic variation, whole-genome resequencing was performed on the contrasting parents. A total of 6.2 and 5.9 million reads were produced for CSR11 and MI48, respectively. Raw reads were passed through the NGS QC Toolkit (Patel and Jain, 2012). The aligned reads were sorted, and putative duplicates were removed, followed by variant calling and variant annotation. The raw variants were filtered at a read-depth cutoff of 5 to proceed with downstream analysis. A total of 9.3 GB and 8.8 GB of cleaned data were obtained for CSR11 and Mi48, respectively, with 92% of the reads mapping to the reference genome for both genotypes. There were 795,968 homozygous polymorphic and 81,625 heterozygous loci found at read depth >=5 between both parents

### Examination of SNPs and Indels

A comparison was made between the two genotypes, and 1962896 SNPs and 127823 indels were identified. Further SNPs and Indels were extracted from within the QTL region. A total of 1368 and 1410 SNPs and 104 and 144 indels were found for MI48 and CSR11, respectively, within the QTL regions (Fig. 1A and B). The transitions and transversions were extracted, revealing a higher number of transitions in MI48 and, vice versa, higher transversions in CSR11. The ratio of Ts to Tv was 2.82 and 2.26 for MI48 and CSR11, respectively. To find out the SNPs and indels in genes within each QTL, SNPs and indels were analyzed for all the QTLs (Fig. 1C and D).

**Figure 1.**
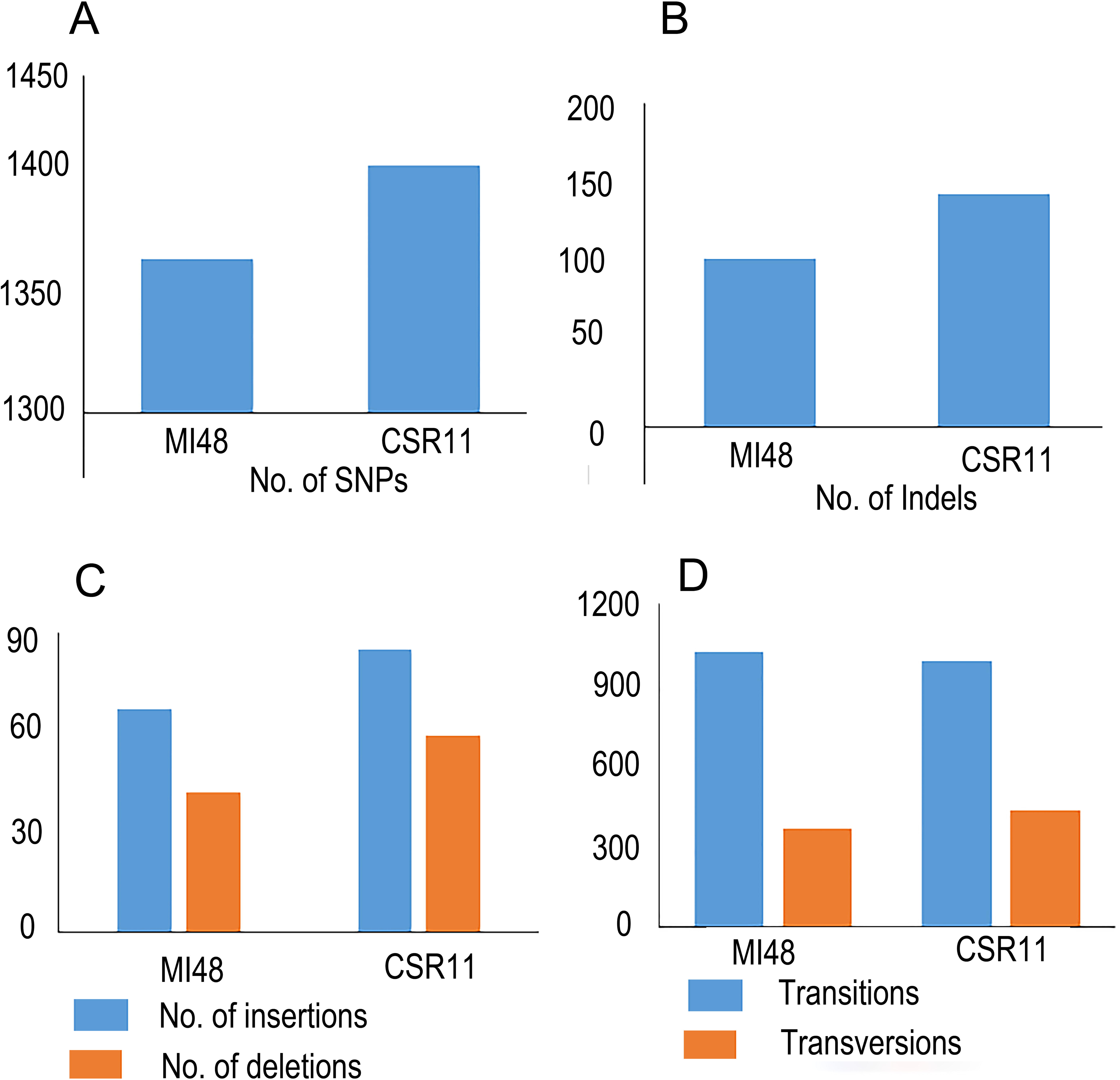
SNPs and Indels allocation in tolerant and sensitive genotypes. (A), SNPs (B), Indels (C), counts of insertion and deletion (D), transitions and transversions.

### Identification and functional annotation of differentially expressed genes (DEG)

Total mRNA from leaves and panicles during the reproductive phase was sequenced from salt-tolerant and salt-sensitive parents. Sequencing total mRNA could improve understanding of the genetic basis of salt tolerance by analyzing gene expression within the associated major QTL region. To analyze differential expression, fold changes were calculated between salt-stress and control conditions in the salt-tolerant (CSR11) and salt-sensitive (MI48) lines. Transcriptome profiling of tolerant and susceptible genotypes from leaf and panicle generates a total of 212102 transcripts (Fig. 2A, B). Cyc and Reactome pathway analyses annotated genes involved in the cell cycle and quantified pathway enrichment for DEGs (Fig. 2C, D). These transcripts were further normalized to assess expression profiles and categorized as upregulated or downregulated based on a fold-change cutoff of ≥2.0 or ≤-2 and a p-value <0.05.

**Figure 2.**
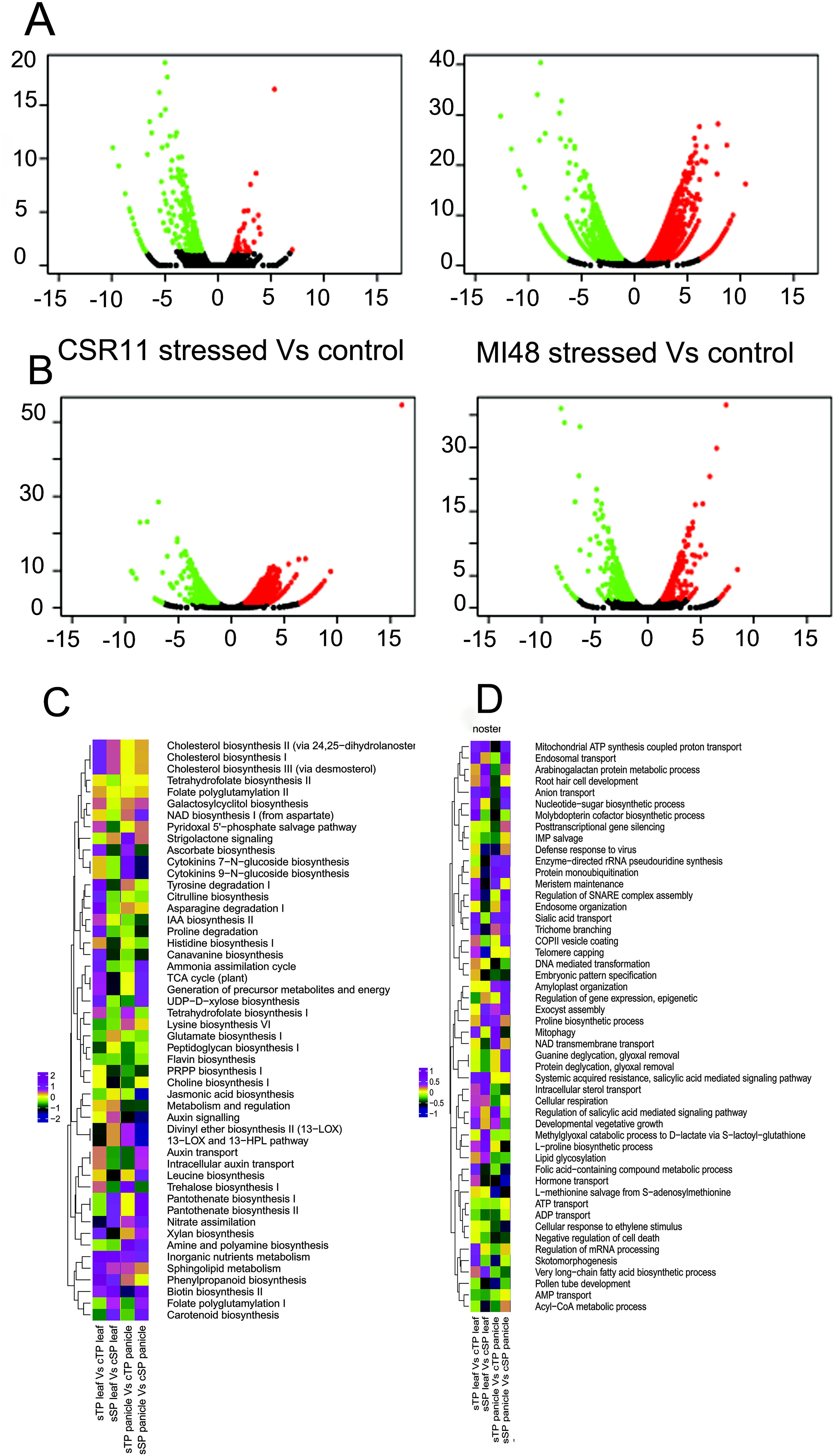
Transcriptome profiling of contrasting rice genotypes under salt stress conditions. (A-B) depicted Volcano plots between control and stressed CSR11 and MI48. Functional enrichment of DEGs showed variation in metabolic pathways (C) and biological and developmental regulations and processes (D), including stress signalling, transport, and energy metabolism.

Finally, a total of 3756 and 1802 DEGs were identified by combining leaf and panicle tissue from susceptible and tolerant genotypes, respectively (Fig. 3). Of these, 1959 (52%) and 283 (15%) transcripts were expressed in leaf tissues of sensitive and tolerant genotypes, respectively. Out of them, 1715 (46%) and 1365 (36%) transcripts were down-regulated and up-regulated, respectively, in the susceptible leaf tissue. In comparison, 729 (40%) and 1073 (60%) transcripts were down- and up-regulated in the tolerant leaf tissue (Fig. 3A-B). A total of 175 (19%) transcripts were down-regulated, and 129 (15%) were up-regulated in susceptible panicles. In comparison, 471 (26%) and 1033 (57%) transcripts were down-regulated and up-regulated, respectively, in panicle tissue (Fig. 3C-D). Since a large number of QTLs have been identified, and to avoid the cumbersomeness of major-effect QTLs showing ≥15% phenotypic variance, an integration of transcriptomics data. The mapping of DEGs over QTLs with the phenotypic variance between 15% and 32% showed that genes on QTLs located on chromosomes (Chr 1, Chr 2, Chr 3, Chr 5, Chr 6, Chr 7, and Chr 8) were differentially regulated under salt stress during the reproductive stage (Fig. 4).

**Figure 3.**
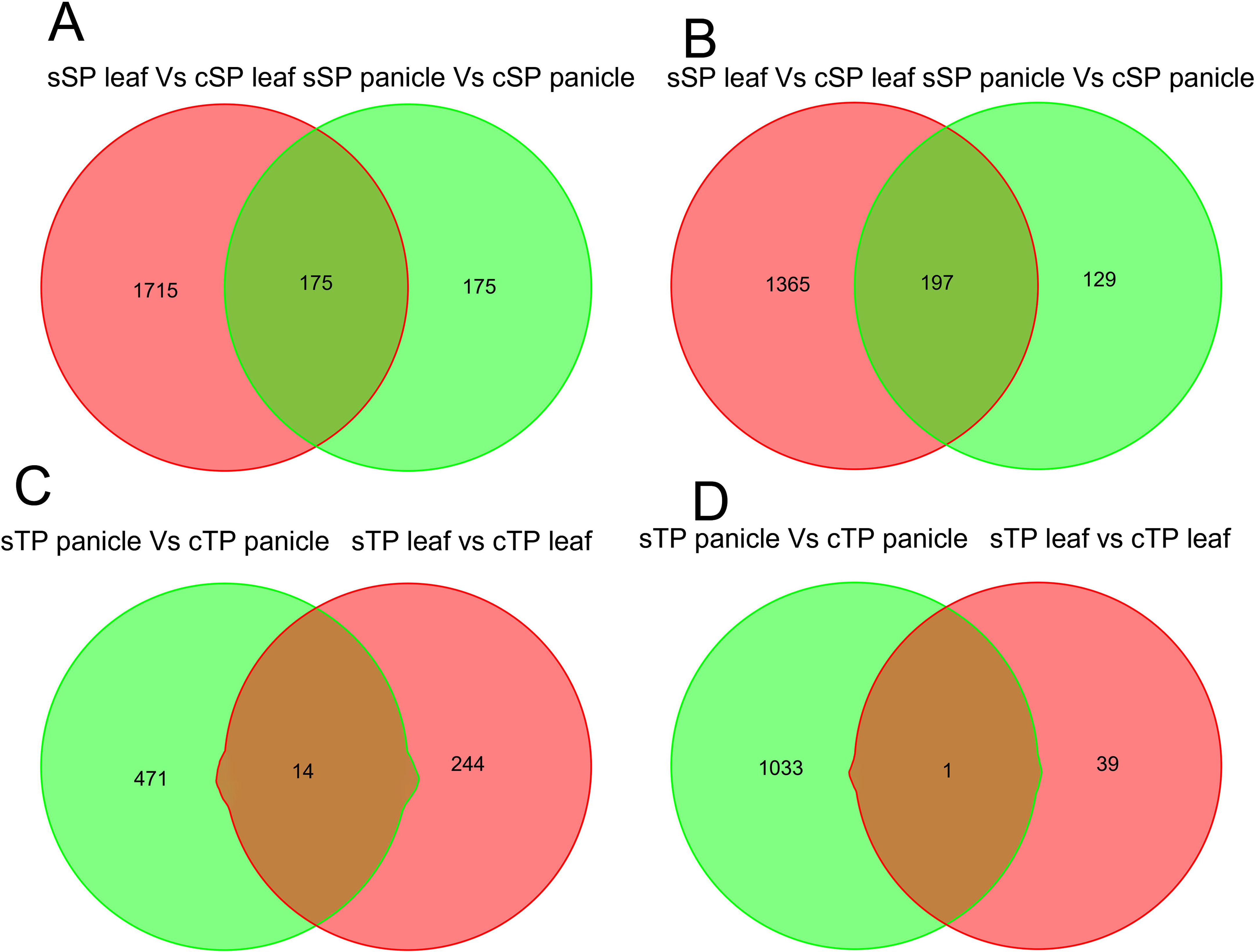
Venn diagram showing DEGs in leaf and panicle tissues of contrasting rice genotypes. The down- and up-regulated genes were shown across multiple comparisons. (A) and (C) Down-regulated DEGs in leaf and panicle tissues (B) and (D) Up-regulated DEGs in leaf and panicle tissues. The figure highlights tissue and genotype-specific transcriptional alterations under salt stress conditions. cSP-control sensitive parent, sSP-salt stressed sensitive parent, cTP-control tolerant parent, sTP-salt stressed tolerant parent.

**Figure 4.**
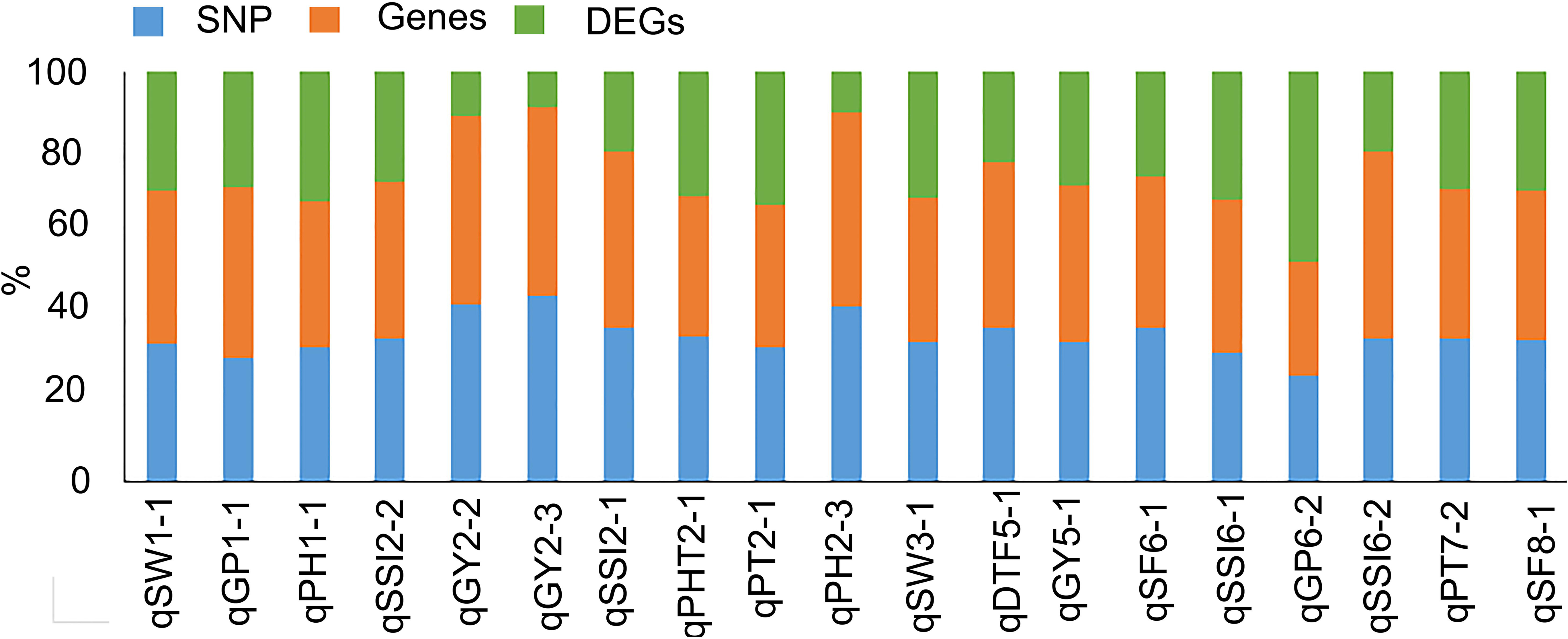
A comparative distribution of genes, SNP variations and DEGs within QTL regions. The stacked bars represent the percentage of SNPs among total genes, along with the total DEGs present in each QTL interval, providing insights into the underlying genomic regions of the genetic and transcriptional architecture.

### QTL for plant height and grain traits are co-localized at chromosome 1

Two major-effect QTLs responsible for different agronomic traits were found at a similar genetic position (17.51 cM) on Chr 1. *qPH1-1* for plant height expressed 19%, whereas *qGP1-1* (grain traits) explained 24% of phenotypic variance, depicting a strong and stable effect of QTLs for both loci (Fig. 5). The left flanking marker was shared by both QTLs (SCR100-Os01g03680); however, the right markers differed and partially overlapped. In the QTL for grains per panicle (*qGP1-1*), 29 genes were differentially expressed in the leaves and panicles of tolerant and sensitive genotypes. Among them, two were multicopper oxidases involved in maintaining Pi homeostasis (*OsLPR3* and *OsLPR5*). In the QTL for plant height (*qPH1*) a total of 267 genes (Supplementary Table 1) showed differential expression, which included different transcription factors like Myb, which helped in the regulation of root growth and coleoptile elongation, and ethylene response (*OsMHZ4*), basic helix-loop-helix (*bHLH*) which mediate seed germination, and also seedling recovery from salt stress (*OsbHLH035*), dehydration and salt stress tolerance DRE binding protein 2 (*OsDREB2A*) TF which helped regulation of the antioxidant response in response to abiotic stresses affected plant growth and development (*OsKEAP1*). There were other genes and transporters were present such as cation efflux protein (*OsMTP9*), zinc fingers, hexose transporter (*OsPGLCT*), universal stress protein, calmodulin-binding protein (CBP), which was responsible for calcium signalling and promotion of root growth under drought stress (*OsCML16*), heat shock proteins 3C (*OsHSPs*), salt-induced AAA-Type ATPase vacuolar sorting protein (*OsSKD1*), peptide transporter for long-distance translocation of dimethylarsinate to rice grain (*OsPTR7, OsNPF8.1*), Skp2-like protein (*OsFBX1*), histone (H2B.1), S-domain receptor-like kinase-11which response to drought and chilling stress in tolerant genotypes, PUB44-interacting protein 1 which negatively regulate disease resistance through WRKY45 (*OsPBI1*), ATP-dependent RNA helicase (*OsRH18*), brassinosteroid insensitive 1-associated receptor kinase 1 precursor (*OsSERL5*), DNA/RNA helicase, DEAD/DEAH (*OsRH1*).

**Figure 5.**
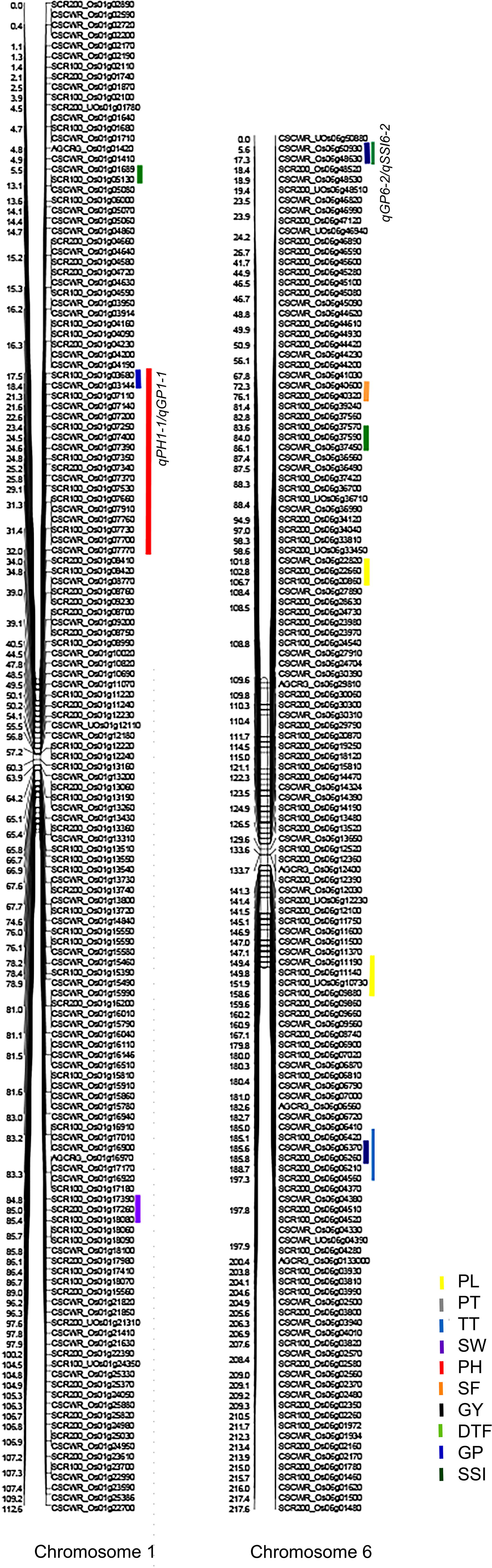
Linkage maps and QTL distribution for yield-related traits in rice. The marker positions and genomic regions were illustrated for chromosomes 1 and 6. Colored bars indicate genomic regions associated with traits. Significant QTL hotspots are highlighted.

Resequencing data were integrated with transcriptomics data for this region to identify candidate genes responsible for the phenotypic effects. The short *qGP1-1* interval was used as the primary region for potential gene analysis, whereas the larger *qPH1-1* interval captured regulatory components of plant height. A total of 10 genes differentially expressed between tolerant and sensitive panicles and leaves had SNPs and indels when compared with CSR11 and MI48 for *qGP1-1* (Supplementary Table 2). Those genes were beta-glucuronosyltransferase GlcAT14A, 3 ubiquitin-protein ligase SINA-like 5, Bowman-Birk type I proteinase inhibitor, and multicopper oxidase LPR1. For *qPH1-1* transcription factor MYB62, plastidic glucose transporter (pGlcT1), NRT1/ PTR7, G-type lectin S-receptor-like serine/threonine-protein kinase, zinc finger protein NFXL2, and NSP-interacting kinase 3 (Supplementary Table 2). In tolerant and sensitive genotypes, 34 and 31 harboured missense variants, and 29 and 27 harboured synonymous variants (Supplementary Table 2). Among them, a missense mutation was detected in Os01g0133400 (Os-pGlcT1), resulting in a change at position 120 from Arginine (R) to Tryptophan (W). This mutation can alter protein folding and signalling in response to salt stress.

### Integration of transcriptomic and sequence variations for the *qGP6-2* / *qSSI6-2* interval

A consistent QTL was identified on Chr 6 at 17.31 cM for two consecutive years under high sodic stress. *qGP6-2* for grain traits and *qSSI6-2* for salt stress index were identified at the same chromosomal positions, and *qSSI6-2* accounted for 32% pf phenotypic variance, suggesting its genetic contribution towards salinity tolerance under stress conditions (Fig. 5). A total of 153 genes were differentially expressed (Supplementary Table 2), which included transcription factor, development of lodicules and stamens (*OsMADS16*), auxin-responsive SAUR gene family member, plant hormone signal transduction, regulation of tiller development (*OsSAUR27*), auxin response factor 19, control of leaf angle (*OsARF7a, OsARF19*), squamosa promoter-binding protein 12 (*OsSPL12*), MYB-CC domain-containing transcription factor, positive regulation of Pi-starvation signaling and Pi-homeostasis (*OsPHR4*), CCCH-type zinc finger protein, negative regulator of the brassinosteroid (BR) response, control of rice architecture via BR signaling (*OsC3H46*). Genes for transporters were P-type heavy metal ATPase, delivery of Zinc to developing tissues (*OsHMA2*), monovalent cation transporter, Na^+^ and K^+^ transport (*OsHKT2;*1; *OsHKT2;4*), peptide transporter/ nitrate transporter (*OsPTR2/OsNRT1*), nitrogen utilization efficiency, growth, and grain yield (*OsPTR9, OsNPF8.2*). Others were bifunctional inhibitor/plant lipid transfer protein/seed storage protein. (*OsLTPG24*), pectin methyltransferase, pistil development during reproductive growth (*OsSTA181, OsPMT16*), Qb-SNARE (soluble N-ethylmaleimide-sensitive factor attachment protein receptor) protein, abiotic stress response (*OsNPSN11*), senescence-associated family protein, basic leucine zipper (bZIP)-containing transcription factor, leaf development (*OsbZIP55*), Raf-Like MAPKKK, mechanical tissue formation at the leaf lamina joint (*OsMAPKKK43*). Further resequencing of the parental lines identified sequence polymorphisms and indels between the tolerant and sensitive lines. A total of 6 missense, 3 synonymous and 7 missense and 1 synonymous SNP variations were detected for the *qGP6-2*/*qSSI6-2* interval (Supplementary Table 2). Therefore, the resequencing and RNA sequencing data were integrated to narrow down the list of potential genes. The nine genes that showed parental polymorphism were differentially expressed in leaves and panicles of tolerant and sensitive rice under sodic stress conditions. They were ANTH/ENTH domain-containing protein (ANTH), subtilisin-like protease SBT2.6, cadmium/zinc-transporting ATPase HMA2, cation transporter HKT9, cation transporter HKT1, auxin-induced protein 10A5, auxin response factor 19, serine/threonine-protein kinase PBL17, protein phosphate starvation response 3 and mitogen-activated protein kinase 12 (Supplementary Table 2). SNP variations in *OsHKT2;1* (Os06g0701700) showed two deletions in the tolerant genotype, one of which was in the coding region of the gene, which was near the C-terminal region explained 3’ regulatory polymorphism. Whereas, *OsHKT2;4* (Os06g0701600) sequence analysis indicated two nucleotide deletions in the tolerant genotype, which were not in the coding region, revealed the presence of a regulatory allele in the tolerant rice genotype. Sequence analysis of *OsANTH12* (Os06g0699800) detected a single SNP variation in tolerant and susceptible lines, which leads to a change in amino acid from Glycine to Arginine at 517 position. Similarly, analysis of *OsPTR2* (Os06g0706400) revealed three SNP variants between tolerant and sensitive lines; among them, two deletions within the coding region were found, which introduced a frameshift mutation in the sensitive line.

## Discussion

Soil salinity and sodicity adversely affect rice growth and yield. To mitigate yield losses, suitable sodicity-tolerant rice cultivars can be developed through marker-assisted breeding (MAB) across diverse genetic backgrounds (Singh et al., 2026; Geetha et al., 2017; Bhandari et al., 2019; Krishnamurthy et al., 2020; Yadav et al., 2020). This study identified and validated novel QTLs associated with salinity stress tolerance at the reproductive stage using a 216-RIL population derived from CSR11 and MI48 (Tiwari et al., 2016). Population genotyping was performed using a 50k SNP array (Singh et al., 2015). Total 19 QTLs contributing ≤ 15% of phenotypic variance were selected for in-depth analysis. Recently, in two BC_1_F_2_ populations derived from Hasawi (salt-tolerant) and BRRI dhan28 (salt sensitive) and CSR28 (salt tolerant) and BRRI dhan 28, using skim sequencing and KASP (Kompetitive allele-specific PCR) technology, 24 and 15 QTLs were discovered for reproductive stage salt tolerance (Mondal et al., 2022). In a BC_1_F_2_ population derived from CSR28 and BRRI dhan 28, three QTLs for *qNF10.1* (number of filled spikelets), *qPFS10.1* (percent of filled spikelets), and *qGY10.1* (grain yield) at the reproductive stage were identified for salinity stress tolerance (Mondal et al., 2025). In the present study, two QTLs of plant height (*qPH1-1*) and grain per panicle (*qGP1-1*) were co-localized at a similar genetic position, suggesting that the chromosomal 1 region may be a QTL hotspot that influences several agronomic traits under salt stress conditions. Besides, these QTLs were identified at the same chromosomal location across years, which demonstrated the stability of QTL under high salt stress conditions. The differences in phenotypic variance across years demonstrated trait-specific tolerance to salt stress, variation in environmental conditions, and stress intensity (Tiwari et al., 2024; Krishnamurthy et al., 2021), rather than inconsistency in QTL identification. The overlap of QTLs suggested either a common gene or a regulatory element, or tight linkage of functionally distinct loci affecting two traits independently. In a reciprocal population derived from Horkuch (salt-tolerant land race) and high-yielding IR29 (salt-sensitive) rice, co-localized QTL at chromosome 1 (175.22 cM) for plant height (at reproductive stage), and shoot length (seedling stage) were identified, which had a positive allele from the Horkuch parent (Haque et al., 2022). In a RIL population developed using Liangxiang5 and 03GY28, a QTL hotspot was detected on chromosome 9, where qleafColor9.1 (19% PV) was mapped and colocalized with *OsSGR,* a senescence-related gene under salt stress conditions (Zhang et al., 2026). In another study using an F_2_ mapping population derived from IR58025A and Giza178 to identify fertility restorer genes and QTLs, three QTLs responsible for pollen fertility were colocalized on chromosome 10, which harbours several loci for pollen fertility with significant phenotypic variance (El-Namaky et al., 2026). Both of the colocalized QTLs in the present study suggested the presence of robust QTLs on chromosomes 1 and 6. Further integration of transcriptomics and sequencing-based analyses was performed to identify the genes underlying QTL regions. Gene expression profiling is a significant approach for analyzing trait-linked genes (Lamb et al., 2006) and helps narrow the number of candidate genes. RNA sequencing was used to analyze developmental responses in panicles and pollen fertility across rice varieties under salt stress, and the KEGG pathway analysis identified significant enrichment of genes involved in flavonoid and phenylpropanoid biosynthesis and protein processing (Duan et al., 2024). In a study of anaerobic germination in rice, differential gene expression profiling was done, target genetic loci were mapped, and 84 candidate genes were identified by combining QTL mapping and transcriptome sequencing (Yang et al., 2019). A similar approach was also used to assess seedling vigour in direct-seeded rice (Sandhu et al., 2023), employing next-generation sequencing (NGS) and transcriptomics. Genes for Pi-homeostasis *OsLPR3* and *OsLPR5*, *OsDREB2A*, peptide transporter for long-distance translocation (*OsPTR7*/ *OsNPF8.1*), calmodulin-binding protein (CBP), which was responsible for calcium signalling and promotion of root growth under drought stress (*OsCML16*), were present under the colocalized QTLs *qPH1-1* and *qGP1-1*. In a reciprocal population of Horkuch and IR29, the reproductive-stage QTL *qPH.1* was enriched for ion-transporter, pollination, and reproductive-process genes (Haque et al., 2022). In the present study, *OsLPR3* and 5 showed higher expression under Pi-deficiency conditions, which played a pivotal role in Pi homeostasis (Cao et al., 2016). Pi deficiency under salinity stress conditions triggered the higher expression of *OsLPR3* and 5. *OsNPF8.1* expression was higher under drought, salinity and N-deficient conditions; the mutant osnpf8.1 had lower grain yield under N-deficient conditions (Diyang et al., 2023), suggesting its role in grain yield. Higher expression of *OsCML16* has been reported under drought, cold, and salt stress in rice (Yang et al., 2020), consistent with the present study, in which its expression was higher under salt stress.

Significant genes that were differentially expressed within the colocalized *qGP6-2* and *qSSI6-2* were *OsANTH12*, subtilisin-like protease (*OsSBT2.6*), control of leaf angle (*OsARF7a, OsARF19*), positive regulation of Pi-starvation signaling and Pi-homeostasis (*OsPHR4*), monovalent cation transporter, Na^+^ and K^+^ transport (*OsHKT2;1, OsHKT2;4*), peptide transporter/ nitrate transporter (*OsPTR2/OsNRT1*), nitrogen utilization efficiency, growth, and grain yield (*OsPTR9, OsNPF8.2*). It was reported that *OsSBT2.6* is involved in cell wall structure and negatively regulates cold tolerance in rice, as the sub4 mutant exhibited increased cold tolerance (Liu et al., 2025). *OsARF19*/*OsARF7a* was involved in floral development, and its overexpression leads to an increase in auxin content, shrunken grains and also enhanced the *OsMADS29* and *OsMADS22* expression signal, which were regulators for floral organ development (Chen et al., 2020). *OsPHR4* was involved in the regulation of Pi-starvation signal transduction and homeostasis (Ruan et al., 2017). In *Eucalyptus grandis, EgPHR* promoter analysis revealed a regulatory transcription factor network, including MYBs and ERFs, which were primarily associated with hormonal responses and abiotic stress (Xu et al., 2025).

*OsHKT2;1* and *OsHKT2;4*, high-affinity plasma membrane K^+^ transporters, are essential for maintaining the ionic balance of Na+ and K+ in plants, especially under K+-deficient conditions (Wang et al., 2024). K^+^ depletion condition induced the expression of *OsHKT2;1* and influx of Na^+^ in the roots (Horie et al., 2007). *OsHKT2;4* was exceptional and showed affinity with not only Na^+^, K^+^, but also to Ca^+2^ and Mg^2+^ to facilitate Mg^+2^ homeostasis in *Xenopus laevis* oocytes (Horie et al., 2011; Zhang et al., 2017), and plants using MGT-type transporters. It was reported that cation/H^+^ (CHXs) and K^+^(Na^+^)/H^+^ (NHXs) transporters maintain pH of the lumen and ionic homeostasis, and Arabidopsis CHXs are expressed in the male gametophyte (Padmanaban et al. 2017; Sze and Chanroj, 2018). Mutants of CHXs could not direct pollen tubes to enter the micropylar ovules (Lu et al., 2011). The most important cation, i.e., K^+^, maintains turgor pressure for pollen tube growth; the K^+^ homeostasis has been maintained through K^+^ channels, cation transporters, and high-affinity K^+^ transporters (Li et al., 2018). *OsHAK26* was highly expressed in anthers and localized to the Golgi apparatus, whereas *OsHAK1* was localized to the plasma membrane. Knockout lines of *OsHAK5* resulted in distorted anthers and reduced pollen germination, indicating its role in pollen viability and grain yield (Li et al., 2022). *OsHKT2;1* and *OsHKT2;4* showed a functional sequence polymorphism that may affect transporter activity through altered gene expression, potentially contributing to K+ homeostasis during salt stress in the male gametophyte and influencing pollen tube growth. Mutant of *AtHKT1;1* accumulated higher Na^+^ in the stamens, leading to male sterility, and overexpression resulted in higher seed yield under salinity stress (Uchiyama et al., 2023). The expression of *OsPTR2* was reported during the grain-filling stages of rice, from 2 to 29 days after pollination, and was higher at the early grain-filling stages than at later stages (Quyang et al., 2010). Higher expression of *OsPTR2* under salt and drought stress conditions, suggesting involvement of *OsPTR2* in the grain filling under salt stress conditions. It was reported that *OsPTR9* overexpression facilitated lateral root development and increased grain yield by improving nitrogen-use efficiency. It mediated the transport of N from the leaf to the panicle, as evidenced by quantifying carbon and nitrogen in mature seeds (Fang et al., 2010) and by agronomic data from field trials of the overexpression lines. Overall, we identified potential candidate genes for reproductive-stage salt tolerance: one on chromosome 1, *Os-pGlcT1* (Os01g0133400), and four on chromosome 6, *OsHKT2;1* (Os06g0701600) and *OsHKT2;4* (Os06g0701700), *OsANTH12* (Os06g0699800) and *OsPTR2* (Os06g0706400). For plant development and adaptation to unfavourable environmental conditions, intracellular compartmentation of sugars is essential. *Os-pGlcT1* protein facilitates glucose transport and is localized in the chloroplast. Overexpression and deletion of the pGlcT1 gene hampered growth under long-day and continuous-light conditions, suggesting its role in starch mobilization (Valifard et al., 2023). A recent study identified *Pv-pGLcT* from *Phaseolus vulgaris* and characterized its role in ribose, fructose, and glucose transport using uptake experiments in *E. coli* (Vob et al., 2025). In addition, *Pv-pGlcT* was also involved in ribose recycling, which is essential for allantoin production. Allantoin-mediated tolerance to salinity was reported via abscisic acid and brassinosteroid pathways in rice and Arabidopsis (Chowrasia et al., 2023; Rajkumari et al., 2023). Pollen tube elongation is critical for pollen germination, as it delivers sperm for double fertilization and sequesters the material via endocytosis. AP180N-terminal homology (ANTH) proteins regulate clathrin-mediated endocytosis. In rice, 17ANTH domain-containing proteins were identified, and *OsANTH3-*driven endocytosis is critical for pollen germination (Lee et al., 2021). In Arabidopsis double mutants for the ANTH-containing protein PICALM5b, pollen tube growth was hampered, thereby facilitating ANXUR protein recycling (Muro et al., 2018). The present study identified the *OsANTH12* gene, which may play an important role in pollen germination under salt stress. Together, the integrative analysis identified the qPH1-1/qGP1-1 and qGP6-2/qSSI6-2 regions as a robust genomic locus for salt stress tolerance, supported by genetic mapping, RNA sequencing, and SNP variation. Candidate genes within the interval should be targeted for functional validation and for marker-assisted development of salt tolerance in rice.

## Data Availability

The RNA sequencing and resequencing data are available in the NCBI Short Read Archive under accession numbers xxxx and xxxx.

## Supplementary Information

The supporting data is available as supplementary figures and tables for the article.

## Acknowledgement

NK, VR and SLK acknowledge funding support from the Indian Council of Agriculture, the National Project on Transgenics in Crops Research (NPTC-3001-2001), and the Department of Biotechnology (BT/PR30273/AGIII/103/1089/2018).

## Author contribution

NK, BPS, AM, AS, NG, ST, and SR helped in experimentation under field and lab conditions. Anoop, PM, and MR did data analysis. PCS, SLK, TT, and SM helped in editing. NKS, VR and SK conceptualized and wrote the whole manuscript.

## Notes

### Competing Interest Statement

The authors have declared no competing interest.

