## Supplementary figures and images for "Integration of QTL Mapping, Transcriptomics, and Genome Resequencing Identifies Yield-Associated Genes for Salt Stress in Rice"

### Supplementary File 1

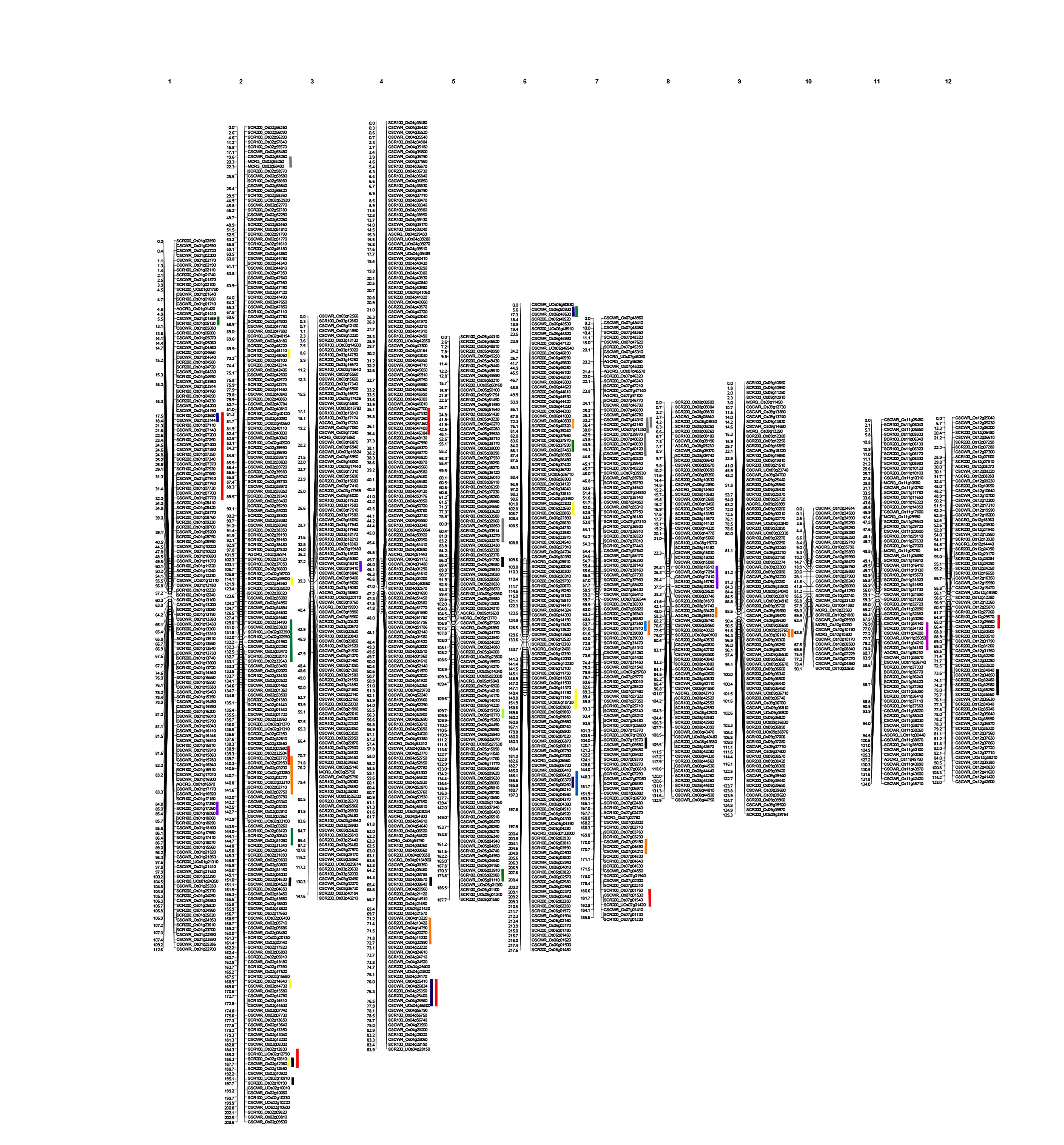
